# Localization and Functional Characterization of the Alternative Oxidase in *Naegleria*

**DOI:** 10.1101/2020.09.26.314807

**Authors:** Diego Cantoni, Ashley Osborne, Najwa Taib, Gary Thompson, Eleanna Kazana, Elizabeth Edrich, Ian R. Brown, Simonetta Gribaldo, Campbell W Gourlay, Anastasios D. Tsaousis

## Abstract

The Alternative oxidase (AOX) is a protein involved in maintaining the Krebs cycle in instances where the respiratory chain has been inhibited, while allowing for the maintenance of cell growth and necessary metabolic processes for survival. Among eukaryotes, alternative oxidases have disperse distribution and are found in plants, fungi and a few protists, including *Naegleria* ssp. *Naegleria* species are free-living unicellular amoeboflagellates, and include the pathogenic species of *N. fowleri*, the so-called brain eating amoeba. Using a multidisciplinary approach, we aimed to understand the evolution, localization and function of AOX and the role that plays in *Naegleria*’s biology. Our analyses suggest that the protein was present in last common ancestor of the genus and structure prediction showed that all functional residues are also present in *Naegleria* species. Using a combination of cellular and biochemical techniques, we also functionally characterize *N. gruberi*’s AOX in its mitochondria and we demonstrate that its inactivation affects its proliferation. Consequently, we discuss the benefits of the presence of this protein in *Naegleria* species, along with its potential pathogenicity role in *N. fowleri*. We predict that our findings will spearhead new explorations to understand the cell biology, metabolism and evolution of *Naegleria* and other free-living relatives.

## INTRODUCTION

*Naegleria gruberi* is a free-living, heterotrophic, microbial eukaryote and a close relative of *N. fowleri*, the so-called “brain-eating amoeba”. It is a non-pathogenic member of the excavate supergroup, which contains key pathogens such as Kinetoplastids (*Trypanosoma, Leishmania*), *Giardia* and *Trichomonas*, and is evolutionarily distant from animals, fungi and plants (Adl *et al*., 2019). *Naegleria* resides primarily as an amoebic (trophozoite) form, but upon environmental stimuli can transform into a flagellate, synthesizing basal bodies and flagella *de novo*, or encyst to allow for dispersion (De Jonckheere *et al*., 2001). It possesses all the major organelles deemed to be canonical for eukaryotes, including nucleus, mitochondria, peroxisomes (Fritz-Laylin *et al*., 2010) and a Golgi (Herman *et al*., 2018). This cellular complexity is reflected in the *N. gruberi* genome sequence (Fritz-Laylin *et al*., 2010) found to encode an extensive complement of cellular machinery and was argued to be reflective of the ancient sophistication present in the last eukaryotic common ancestor (Koonin, 2010).

Found in soils and freshwater worldwide, *N. gruberi* can thrive in a wide range of osmotic and oxygenic conditions (De Jonckheere, 1979, 2014; Tyml *et al*., 2016). It has the capacity for full aerobic and anaerobic metabolism, and was predicted to have assimilated unique biochemical adaptations both within and outside the mitochondria (Fritz-Laylin *et al*., 2010, 2011; Ginger *et al*., 2010). Among those, only a handful of pathways have been localized and characterized, in the trophozoite stage of this microbial eukaryote. Some examples include the cytosolic localization and functional characterization of the [FeFe]-hyderogenase (Tsaousis *et al*., 2014), the mitochondrial localization and functional characterization of ferritin (Mach *et al*., 2018), and the oxygen-depended metabolism of lipids (Bexkens *et al*., 2018). While the last report provided a hint on the function of *Naegleria*’s alternative oxidase (AOX), a thorough investigation on this important oxygen-depended enzyme is lacking.

The alternative oxidase (AOX) is a terminal oxidase typically involved in bypassing the electron transport chain in plant mitochondria, even though it has also been identified and localized in the mitochondria and related organelles of many non-related microbial eukaryotes including trypanosomes (Clarkson *et al*., 1989), *Candida albicans* (Yan *et al*., 2009), *Cryptosporidium* (Roberts *et al*., 2004) and *Blastocystis* (Stechmann *et al*., 2008; Tsaousis *et al*., 2018). Due to its absence in humans, the protein is considered a potential drug target, and has been well studied in some of these pathogenic species (Shiba *et al*., 2013; Tsaousis *et al*., 2018; Duvenage, Munro and Gourlay, 2019). Despite the wide distribution and extensive research on this group of proteins, their overall physiological roles are still unclear (Moore and Albury, 2008). Intriguingly, it has been suggested that AOX may be involved in maintaining tricarboxylic acid cycle turnover under high cytosolic phosphorylation potential, stress tolerance (oxygen), and thermogenesis (Finnegan, Soole and Umbach, 2004; Moore *et al*., 2013).

Herein, we employed a multiphasic approach to characterize the AOX of *N. gruberi*, while comparing it with both the *N. fowleri* and *N. lovaniensis* homologues and examining their origins. Upon structural characterization of all the counterparts, we generated a specific polyclonal antibody against *N. gruberi* AOX, with which we localized the protein in *N. gruberi’s* mitochondria using a combination of assorted cellular approaches. Experiments with high-resolution respirometry demonstrated that the *N. gruberi* homologue confers cyanide resistance. This study represents the first thorough characterization of AOX in *Naegleria* species, which could provide further understanding into the biochemical adaptations of this peculiar and highly adaptable microbial eukaryote.

## MATERIALS AND METHODS

### Bioinformatic Analysis

The predicted amino acid sequences of the alternative oxidase homologues of *N. fowleri* and *N. lovaniensis* were obtained from Genbank (NCBI) using the following accession numbers; GCA_008403515.1 (*N. fowleri*), GCA_003324165.1 (*N. lovaniensis)*. The amino acid sequence of the AOX from *N. gruberi* was obtained from uniprot (D2V4B2). The Phyre2 web portal for protein modelling, prediction and analysis was used in intensive mode (Kelley *et al*., 2015) to derive the structure for *Ng*AOX. Structures were then downloaded and analyzed using PyMol. For sequence alignment Clustal Omega (Sievers *et al*., 2011) was used, output was downloaded as fasta file and analyzed using Jalview 2.11.0 (Waterhouse *et al*., 2009). Sequence identifiers for AOX alignments were: *Trypanosoma brucei brucei* (Q26710), *Cryptosporidium parvum* (Q6W3R4), yeast; *Candida albicans* (A0A1D8PEM4), plant; *Arabidopsis thaliana* (Q39219), and, fungi; *Neurospora crassa* (Q01355).

### Phylogenetic analysis

Three local databanks of proteomes, representative of all diversity from eukaryotes, bacteria and proteobacteria were assembled from the National Center for Biotechnology Information (NCBI): 193 proteomes from Eukaryotes (one per genus); 1017 proteomes from Bacteria (3 per family) and 1082 proteomes from Proteobacteria (1 per genus). Homology searches were performed using HMMSEARCH, from the HMMER-3.1b2 package (Johnson, Eddy and Portugaly, 2010), with the option --cut_ga to screen all the proteomes in the three databanks for the presence of AOX pfam domain (PF01786.17). All the hits were then manually curated in order to discard false positives. The remaining hits were aligned using MAFFT-v7.407 (Katoh and Standley, 2013) with the linsi option and trimmed with BMGE-1.1 (Criscuolo and Gribaldo, 2010) using the BLOSUM30 substitution matrix to select unambiguously aligned positions. A maximum likelihood tree was then generated using IQTREE-1.6.12 (Nguyen *et al*., 2015) under the TEST option with 1000 ultrafast bootstrap replicates.

### Cell Culture

*N. gruberi* strain NEG-M (kindly provided by Lillian Fritz-Laylin, Biology Department, University of Massachusetts, Amherst, USA) was cultured in M7 media at 28 °C (Tsaousis *et al*., 2014). Cells were passaged every 3 to 5 days to prevent overconfluency.

### Antibody generation

A 16 amino acids peptide (nh2-CMHRDYNHDMSDKHRA −conh2) was designed by Eurogentec based on the provide sequence of AOX of *Naegleria gruberi* (XP_002672918) (see **Suppl Figure S2**). 5 mg of the peptide was coupled with the carrier protein KHL (Keyhole Limpet Hemocyanin) and was subsequently used to inoculate a rabbit (through the Eurogentec’s speedy program) for antibody production. The antibody’s affinity (Eurogentec; Peptide: 1911009, Rabbit 237) was confirmed through ELISA.

### Cell Fractionation and western blots

To separate organelles from cytosol, cell fractionation by centrifugation was carried out as previously described (Herman *et al*., 2018). Lysates were mixed with 4x sample buffer and heated to 95 °C for 10 minutes. Samples were then loaded in two tris-glycine gels for gel electrophoresis. One gel was subjected to Coomassie staining overnight and destained the following day to assess equal loading. The other gel was transferred to PVDF membrane using a trans-blot turbo transfer system according to manufacturer’s protocol (Bio-rad). Membranes were blocked with 5% milk in TBS buffer containing 0.5% tween-20 for 1 hour at room temperature. Primary antibody staining was carried out overnight at 4 °C with the following antibody dilutions: anti-AOX 1:1000, and previously published antibodies anti-HydE 1:1000 and, SdhB 1:1000 (Tsaousis *et al*., 2014). Membranes were washed four times with TBS-T for 5 minutes prior to secondary staining with HRP-conjugated goat anti-mouse and goat anti-rabbit antibodies (Invitrogen). For detection, membranes were incubated with ECL reagent (Bio-rad) for 30 seconds and imaged using syngene G:BOX imager. Membranes were stripped with mild stripping buffer composed of 13 mM glycine, 3.5 mM sodium dodecyl sulfate, 1% tween-20, pH 2.2. PVDF membranes were washed twice for 10 minutes in mild stripping buffer, followed by two washes in PBS for 5 minutes, and lastly two washes for 5 minutes with TBS-T, prior to blocking for the next immunoprobe.

### Immunofluorescence Microscopy

*N. gruberi* was seeded onto sterile poly-L-lysine treated glass coverslips in a 6-well plate and incubated overnight at 25 °C. The following day the cells were treated with 250 nM MitoTracker Red for 30 minutes. The media was then aspirated and cells were washed with 1x PBS, followed by fixation using 2% formaldehyde for 20 minutes. After fixation, 2% formaldehyde was removed and cells were permeabilised with 0.1% triton-X 100 for 10 minutes. After permeabilization cells were washed three times with 1x PBS and blocked using 3% bovine serum albumin in PBS for 1 hour. Primary antibody staining was carried out at room temperature for 1 hour with custom made AOX antibodies (Eurogentec; Peptide: 1911009, Rabbit 237), diluted to 1:1000. Secondary antibody staining was carried out for 1 hour in the dark using anti-Rabbit-IgG-Alexa 488. Slides were washed and mounted using Prolong Gold Antifade with DAPI. Laser Confocal Microscopy was carried out using the LSM 880 Laser Confocal with Airyscan by Zeiss. Laser sets used were 405, 488 and 594, with airyscan detector plate imaging for high resolution. Images were captured and analyzed using Zen software suite by Zeiss.

### Immunoelectron Microscopy

Samples were prepared as previously described (Herman *et al*., 2018). Blocking of the samples was achieved via a 1 hour incubation in 2% BSA in PBS with 0.05% Tween 20. Primary antibody staining was performed by incubating AOX antibodies at 1:10, 1:50 and 1:100 dilutions for 15 hours at 8 °C. The sample grids were then incubated for 30 minutes at room temperature, with the corresponding gold-conjugated secondary antibodies. Counter-staining was achieved by incubation with 4.5% uranyl acetate in PBS for 15 minutes and a 2-minute incubation in Reynold’s lead citrate. The sample grids were imaged with a Jeol 1230 Transmission Electron Microscope operated at 80kV and images were captured with a Gatan One view digital camera.

### High Resolution Respirometry

Real time respirometry was monitored using an OROBOROS Oxygraph-2k with Clark polarographic oxygen electrodes and automatic titration-injection micropump. The chambers of the oxygraph were calibrated using 2 ml of M7 media without glucose for 20 minutes. *N. gruberi* cells were seeded in the chamber at a density of 100,000 cells per ml. Respiration was monitored before and during drug additions. The drugs used to assess respiration were added in the following order; 1 mM potassium cyanide, 5 μM antimycin A and 1.5 mM salicylhydroxamic acid (SHAM). The experiment was then repeated with the drug additions in reverse order. Student-t test was used to determine significance.

### Cell Proliferation Assay

*Naegleria* cells were seeded at a density of 2000 cells per well of a clear F-bottom 96 well plate and incubated in identical conditions as stated above. The plate was placed on a Juli™Stage live cell monitoring system, programmed to capture an image of each well every hour for 5 days. On day-2, wells were inoculated with concentrations of SHAM at 1 mM, 0.1 mM, 0.001 mM and, 0.001 mM. Plates were then incubated for a further 3 days. At the end of the experiment the images were used to count cell numbers using ImageJ. Using an image with a scale bar, we used the grid function with a known area per square that was applied to all images taken at 0, 24, 48, 72, 96 and 120 hours. Using the cell counter function, we counted the number of cells in a grid square, and multiplied it to the surface area size of a F-bottom 96-well plate in order to estimate the total cell number per well. The experiment was completed with three biological replicates, whereby each biological replicate consisted of three technical replicates. Cell counts were graphed using GraphPad Prism 8.

## RESULTS

### Phylogenetic analysis of alternative oxidase proteins

We carried out an exhaustive search for AOX homologs in updated eukaryotic and bacterial databanks. As bacterial hits were mainly from proteobacteria, we also searched for AOX homologs in a proteobacteria databank with more diversity within this phylum. We identified 323 AOX homologs: 265 from 193 eukaryotic taxa, 56 from 1,082 proteobacterial taxa, and 2 from 896 bacterial taxa (other than proteobacteria) (**Suppl table S1**).

Regarding the eukaryotic distribution, in addition to previously described groups Alveolata, Euglenozoa, Metazoa, Choanoflagellates, Stramenopiles, Fungi, Rhodophya, Heterolobosea and Viridiplantae (Pennisi *et al*., 2016), we identified AOX homologs in some eukaryotes that were not reported: Apusuzoa, Amoebozoa, Filasterea, Haptophyceae and Rhizaria. No homologs were found in Kipferlia, Metamonada and Hexamitida (**Suppl Figure S1** and **Suppl table S1**). Bacterial hits were much less diversified, as they are restricted to alphaproteobacteria, betaproteobacteria and gammaproteobacteria. The other two hits belong to Bacteroidetes and the CPR (Candidate Phyla Radiation) and they probably correspond to transfers from proteobacterial taxa (**Suppl Figure S1**). Our phylogenetic analysis shows that AOX homologs of Metazoans (UFB=86%), Haptophyceae, Choanoflagellata (UFB=100%) and Heterolobosea (UFB= 100%; inset **Figure 1**) form distinct and monophyletic groups pointing toward the presence of an AOX in the ancestors of each group. Regarding bacterial sequences, although they branch together and form a monophyletic clade (UFB= 97%), it seems that proteobacterial taxa transferred AOX sequences to other bacteria (Bacteroidetes and CPR) and to some Euglenozoa. This bacterial clade forms a sister group of Streptophyta nad Rhodophyta (UFB= 97%), however, it is not clear which eukaryotic group transferred AOX sequences to proteobacteria.

**Figure 1.**
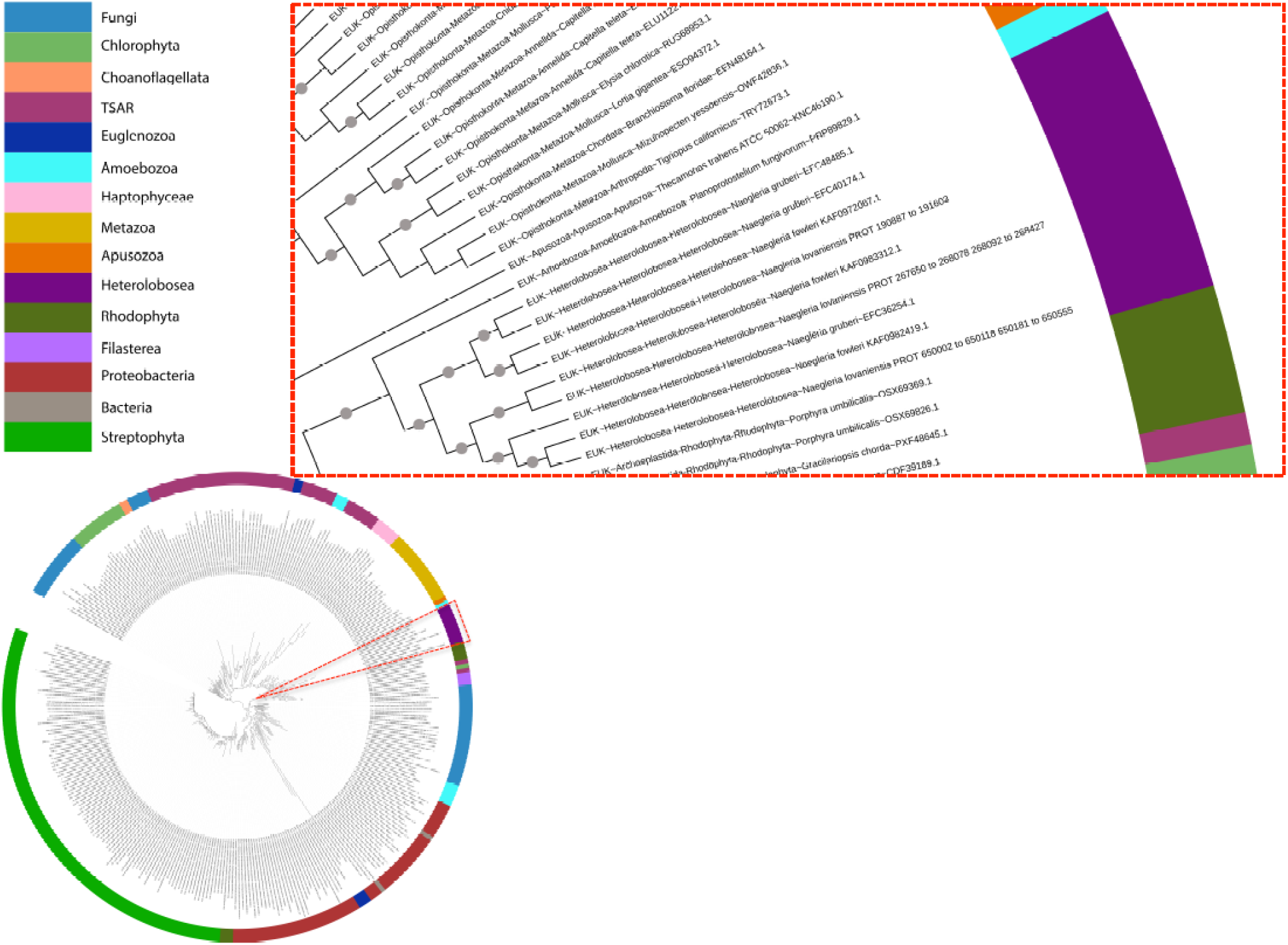
Phylogenetic trees demonstrating the origin of alternative oxidases in heterobolosea. Maximum likelihood tree of AOX homologs in eukaryotes and bacteria (323 sequences, 148 amino acid positions). The tree was inferred with IQTREE using the LG+I+G4 model selected under the BIC criterion. Grey dots correspond to supports higher than 80%. The scale bar corresponds to the average number of substitutions per site. Inset focus on the heterobolsean section of the tree, demonstrating the evolution and duplications events through the *Naegleria* genus. Full phylogenetic tree can be found in **Suppl. Figure S1**.

The presence of the two separate and monophyletic groups Streptophyta (UFB= 100%), Chlorophyta (UFB=82%) shows that, although the presence of AOX could not be inferred in the ancestor of Archaeaplastida, it can be inferred in the ancestors of both groups. Finally, regarding Fungi, the presence of three separate and monophyletic groups containing Ascomycota and Basidiomycota (UFB= 80%), Chytridiomycota (UFB= 100%) and a mix of Zoopagomycota, Chytridiomycota, Cryptomycota, Mucoromycota, Blastocladiomycota and Microsporidia (UFB= 74%), suggests that AOX was acquired at least three times independently in this group.

### Amino acid alignments reveal conserved residues in *Naegleria gruberi, N. fowleri* and *N. lovaniensis*

The amino acid sequences of AOX in *N. gruberi, N. fowleri* and *N. lovaniensis*, were aligned against the AOX found in protists; *Trypanosoma brucei brucei, Cryptosporidium parvum*, yeast; *Candida albicans*, plant; *Arabidopsis thaliana* and, fungus; *Neurospora crassa*. The N-termini displayed little conservation. However, key conserved residues were detected across all species from the middle of the sequences, continuing towards the C-terminus (**Suppl. Figure S2**). Most of the conserved residues localized in the alpha helical arms, including the residues responsible for membrane interaction, diiron binding domain and, three universally conserved tyrosines (**Suppl Table S2**). The presence of these key features strongly suggests that *Naegleria*’s predicted AOX would retain functional activity.

### Structure modelling of AOX from *Naegleria gruberi*

Using Phyre2, we were able to generate models of the *Ng*AOX monomer. The resulting model reveals a monomer containing the characteristic four-bundle helix, in which resides the active site (**Figure 2**). The active site is composed of the characteristic four glutamate residues and two histidine residues which are responsible for diiron binding. In addition, the universally conserved tyrosine residue necessary for activity is present at residue number 175. Furthermore, helices α1 and α4 contains a strong hydrophobic region, also a key feature of AOX, as these helices are involved in membrane insertion. Amino acid conservation of glycine 85 and glycine 165 between all *Naegleria* species and *Trypanosoma* contribute to the structural kink of both α2 and α5 helices. We also observed conservation between amino acids responsible for membrane-binding regions and for dimerization (**Suppl Table S2)**. These amino acid conservations highlight the conserved structure of AOX, its characteristic helices and hydrophobicity patch for membrane anchoring.

**Figure 2.**
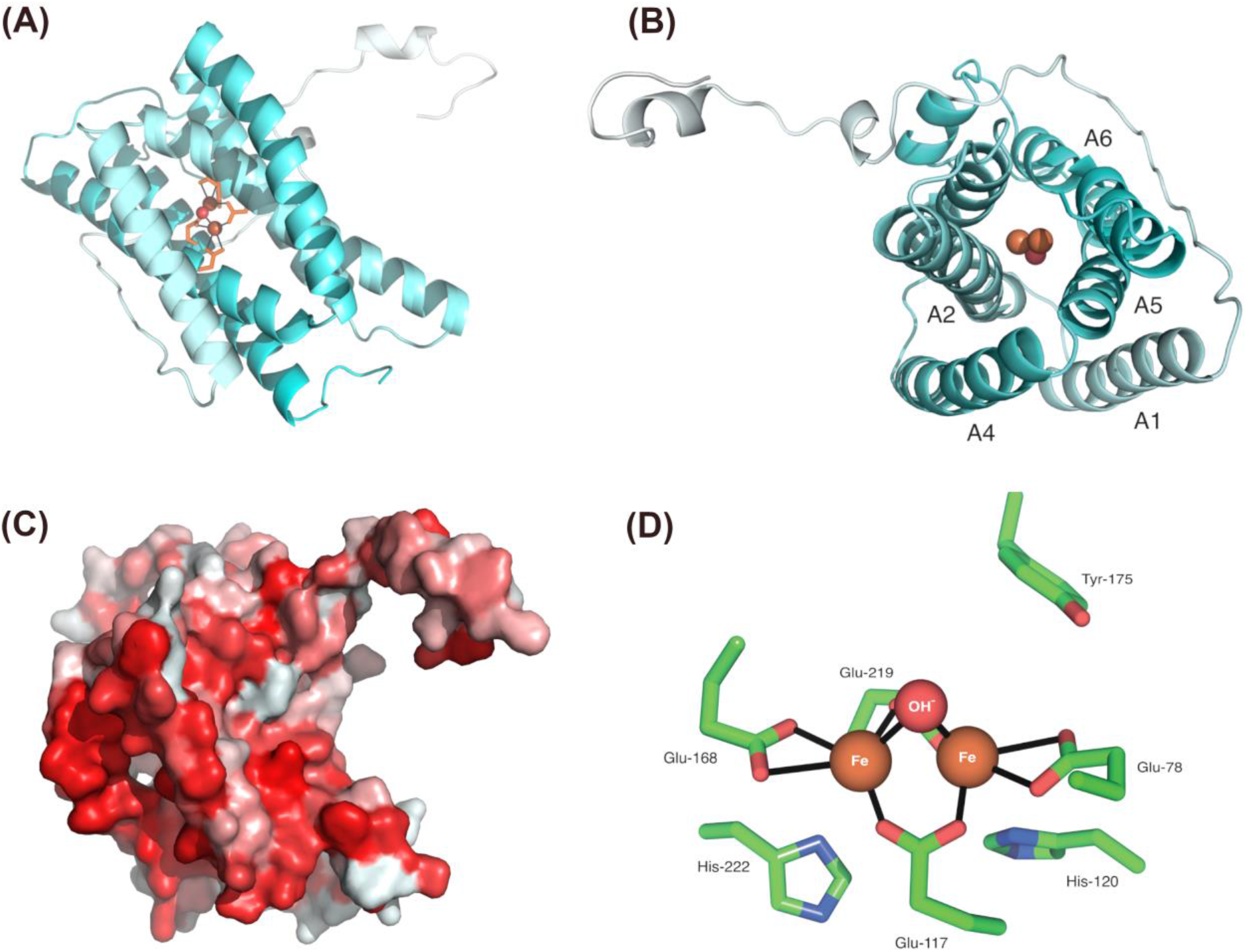
Structural modelling of *N. gruberi* AOX reveals canonical features. Using Phyre2 we modelled the structure of ngAOX. We observed the presence of the di-iron domain and alpha-helical bundles, as viewed from the side (A) and top-down (B). Rendering the structure by hydrophobicity shows the typical hydrophobic patch required for membrane anchoring (C).

### Localization of AOX in *N. gruberi*

To verify the presence and localization of AOX, fractionation by centrifugation was carried out to isolate the mitochondria and cytosol. Using a custom-made antibody raised against *N. gruberi* AOX, our western blot analysis reveals the presence of AOX in the mitochondrial fraction and not in the cytosolic fraction (**Figure 3C**) Successful fractionation was confirmed by immunoblotting Hydrogenase E protein, which localizes exclusively in the cytosol and succinate dehydrogenase, a previously confirmed mitochondrial marker (Tsaousis *et al*., 2014). The localization of AOX was also carried out by immunofluorescent microscopy, whereby *N. gruberi* cells were stained with mitotracker red before fixation, and subsequently fixed and immunostained for detecting AOX. Our imaging reveals a high degree of colocalization between the mitotracker signal and the AOX signal derived from the secondary antibodies. This indicates that AOX is localized in the mitochondria (**Figure 3A & B & Suppl. Figure S3**). To further confirm these observations, we have subsequently carried out immunoelectron microscopy (IEM) on resin fixed *N. gruberi* sample grids. As a result, we detected positive signal inside derived from the immunogold labelling, strongly suggesting its localization is within the inner membrane of the mitochondria (**Figure 4**).

**Figure 3.**
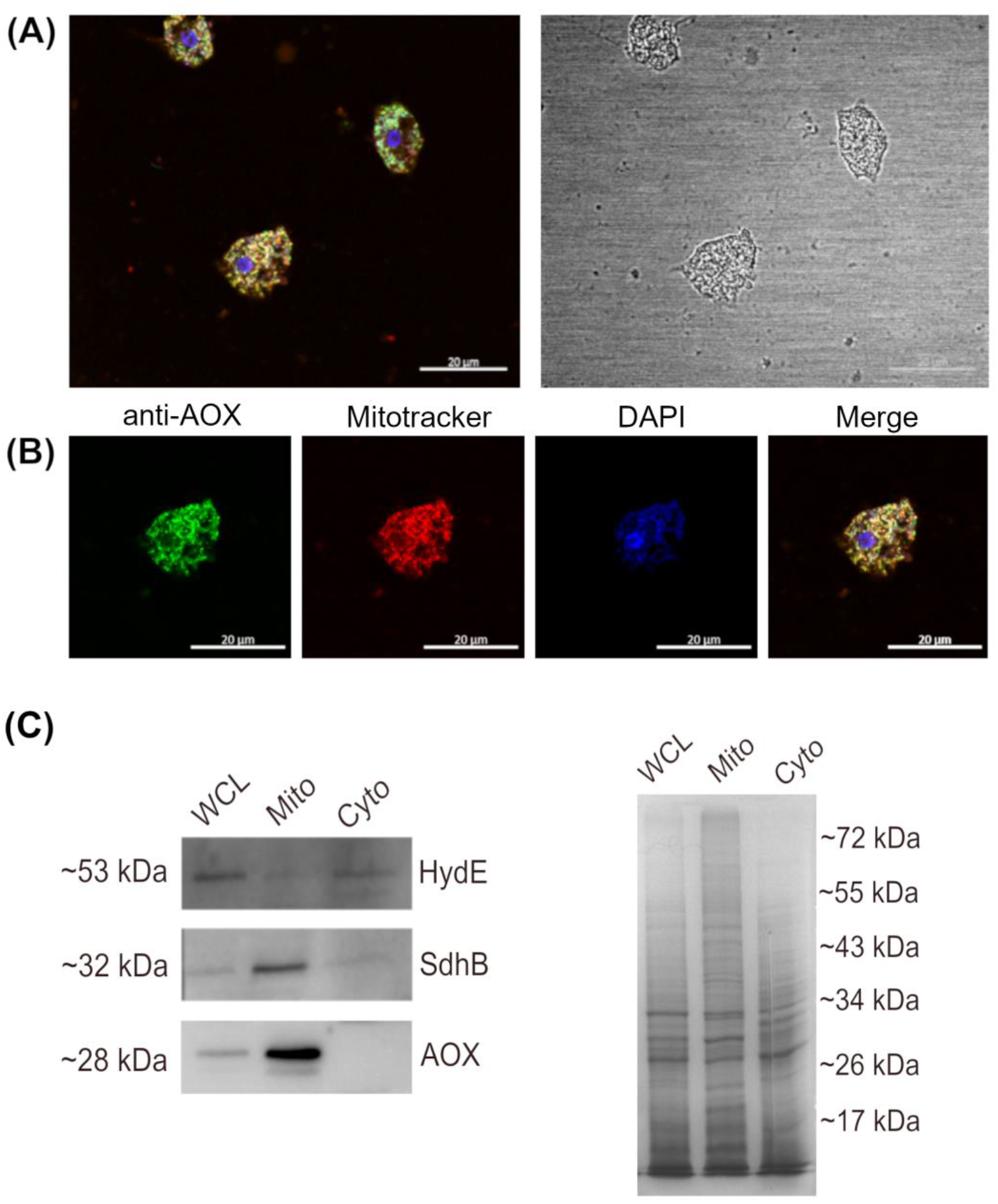
Confocal microscopy and western blot reveals mitochondrial localization of AOX. (**A & B**) *N. gruberi* cells were treated with mitotracker red prior to fixation and then probed with AOX antibodies (green). Nuclear marker and mtDNA staining is shown in blue (DAPI staining) Our confocal imaging reveals a high degree of co-localization between the mitotracker red signal and the green signal derived from immunoprobing AOX. These results provide visual confirmation of the expression of AOX and their localization in the mitochondria. Localization figures of more *N. gruberi* cells can be found in **Suppl. Figure S3**. For western blotting lysates were fractionated by centrifugation to yield samples containing pelleted mitochondrial fraction and a clarified cytosolic fraction. Successful fractionation from the whole cell lysate was confirmed by immunblotting (lefthand side) for hydrogenase maturase E (HydE) that localizes in the cytosol, and succinate dehydrogenase B (SdhB), which localizes exclusively in the mitochondria (**C**). Immunostaining against AOX revealed a band around ~28 kDa in the whole cell lysate and mitochondrial fraction only. A Coomassie stain was carried out to the parallel SDS-PAGE gel parallel to assess equal loading between samples (right-hand side).

**Figure 4.**
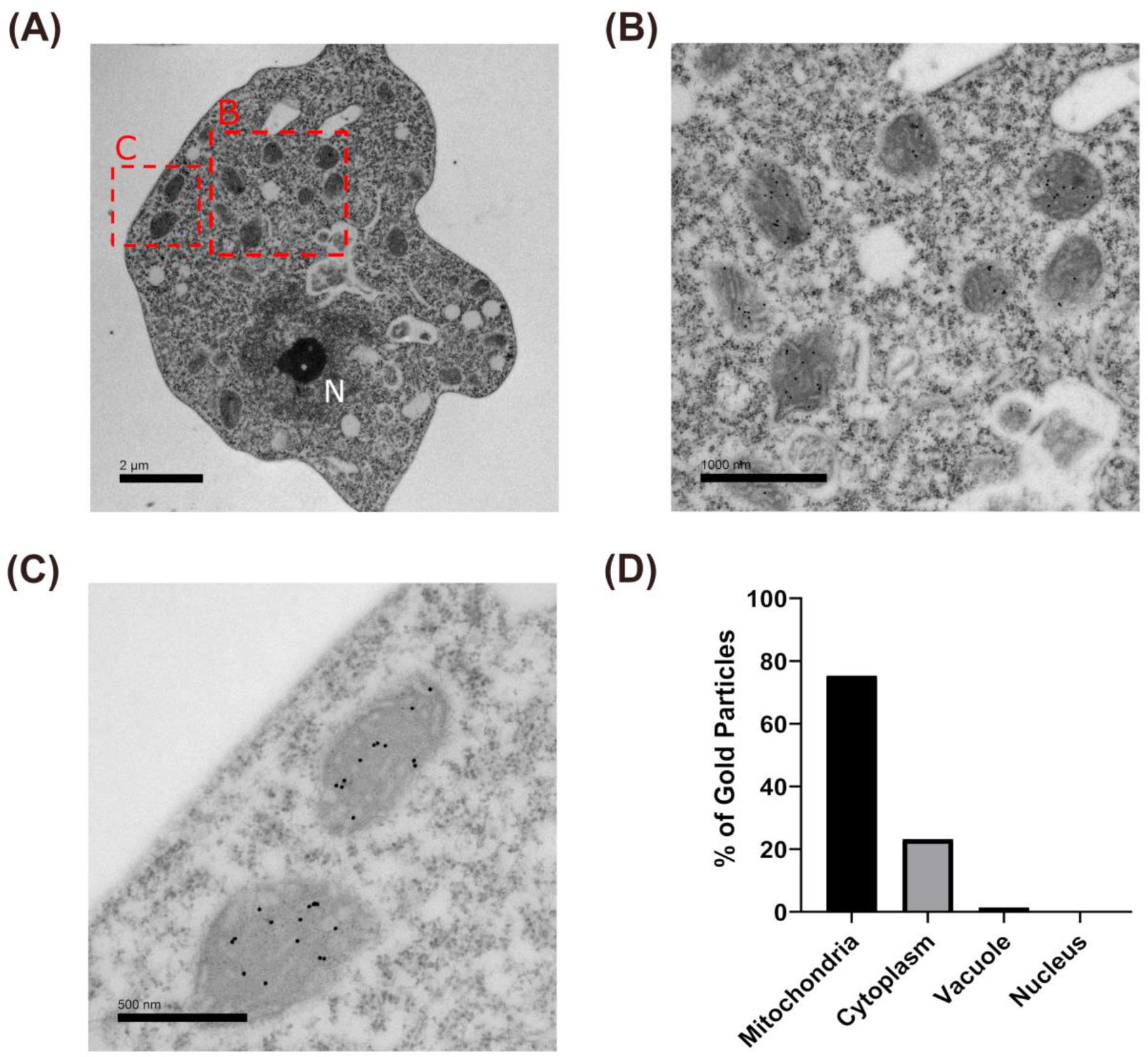
Immunoelectron microscopy reveals AOX localization within the inner mitochondrial membrane. Fixed *N. gruberi* samples were probed for IEM to assess their localization in the mitochondria. The AOX signal derived by the immunogold secondary antibodies bound against AOX primary antibodies localizes predominantly inside the mitochondria.

### Real time respirometry reveals AOX confers cyanide resistance

In order to assess whether *N. gruberi* respires from either the classical respiratory chain or through AOX, real-time respirometry data was collected. Cells were cultured for three days and washed with M7 media without the addition of glucose. Afterwards, 2 ml of cell resuspension was seeded into the respirometer chambers at a density of 10^5^ cells per ml. After establishing the routine respiration, a series of drugs were added sequentially to each chamber and oxygen flux was monitored (**Figure 5**). Initially, 1 mM of complex IV inhibitor potassium cyanide was injected into the chambers, revealing an increase in oxygen flux and decrease in oxygen concentration, which would suggest that respiration via alternative oxidase pathway is active, as the presence of cyanide has no effect on *N. gruberi*. The addition of complex III inhibitor, Antimycin A, had no effect on respiration. Lastly, addition of 1.5 mM of AOX inhibitor SHAM resulted in a significant decrease in respiration (**Figure 5A, B**). When the experiment was repeated with drug additions in the reverse order, the SHAM treatment resulted in a significant decrease in respiration, whereas Antimycin A and KCN had no further effects on respiration (**Figure 5B, C**). This would suggest that *N. gruberi* respire predominantly via the AOX pathway.

**Figure 5.**
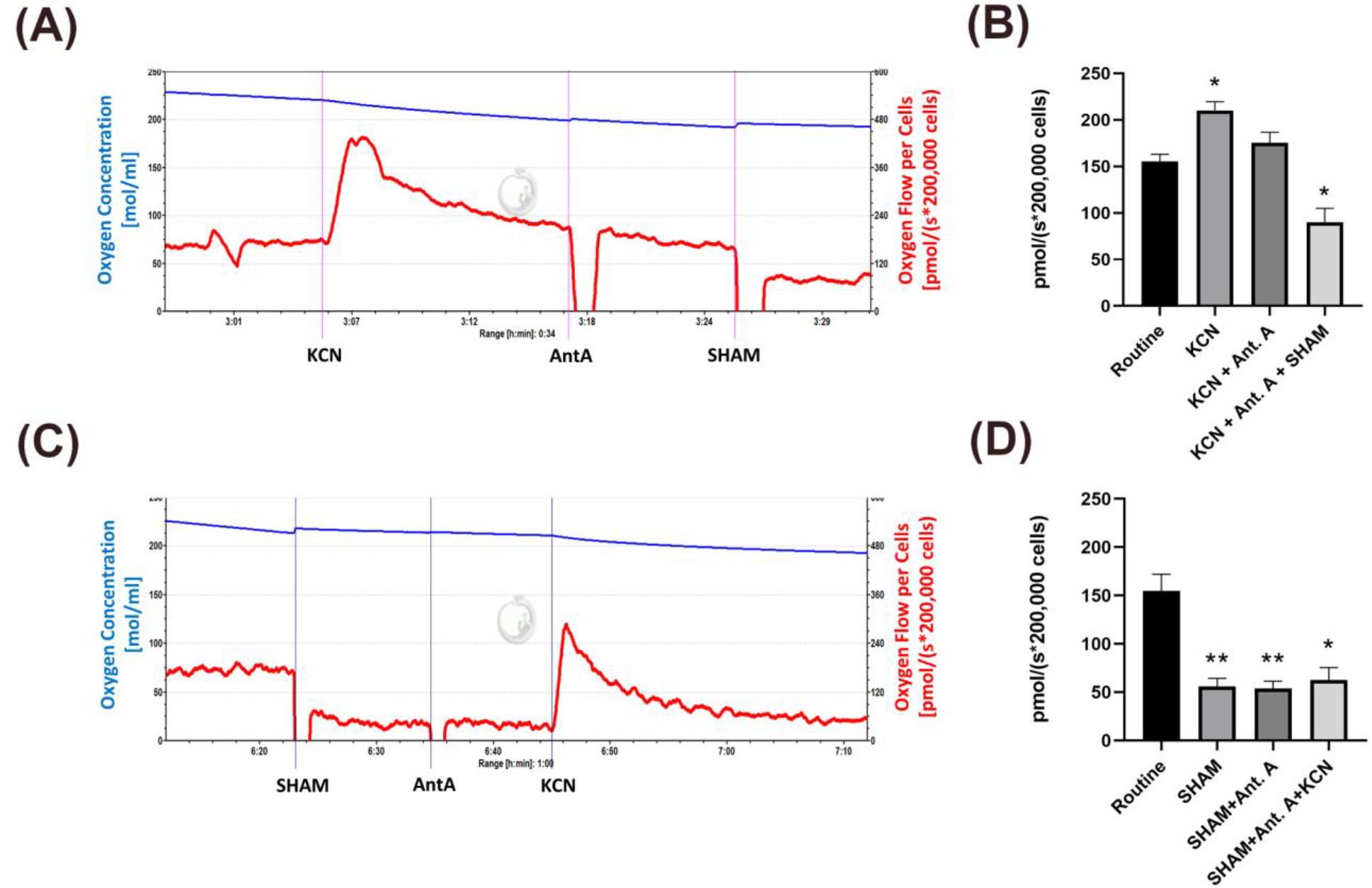
High resolution real-time respirometry reveals cyanide resistant respiration. *N. gruberi* cells were subjected to real-time respirometry in the presence of metabolic inhibitors to assess respiration. Cells were added to the chambers without the presence of inhibitors to assess normal respiration, followed by the addition of a mitochondrial complex inhibitor. The order of inhibitors were KCN (complex IV inhibitor) followed by Antamycin A (complex III inhibitor) and SHAM (AOX inhibitor) (A). The experiments were then completed in reverse order (B). KCN did not display decreases in respiration, whereas SHAM significantly reduced respiration. P values; * <0.05, **<0.01

### The presence of SHAM in culture media reduces cellular proliferation

Due to the decrease respiration in *N. gruberi* in the presence of SHAM, we opted to evaluate cell growth under varying concentrations of SHAM using a Juli™Stage live cell monitoring system (**Figure 6**). Cells were grown in 96-well culture plates for 48 hours to allow proliferation. At the 48 hour mark, we pipetted varying amounts of SHAM in culture media to reach the following final concentrations; 1 mM, 0.1mM, 0.01 mM and, 0.001 mM. The plate was then returned to the incubator for another 72 hours. By counting the number of cells under each condition, we were able to verify the negative effect of SHAM on proliferation. Concentrations of SHAM at 1 mM to 0.1 mM were effective at stopping cell proliferation altogether, whereas 0.01 mM SHAM greatly reduced cell proliferation. SHAM concentration of 0.001 mM had no effect on cell proliferation, as the cell counts followed a similar pattern to the negative control.

**Figure 6.**
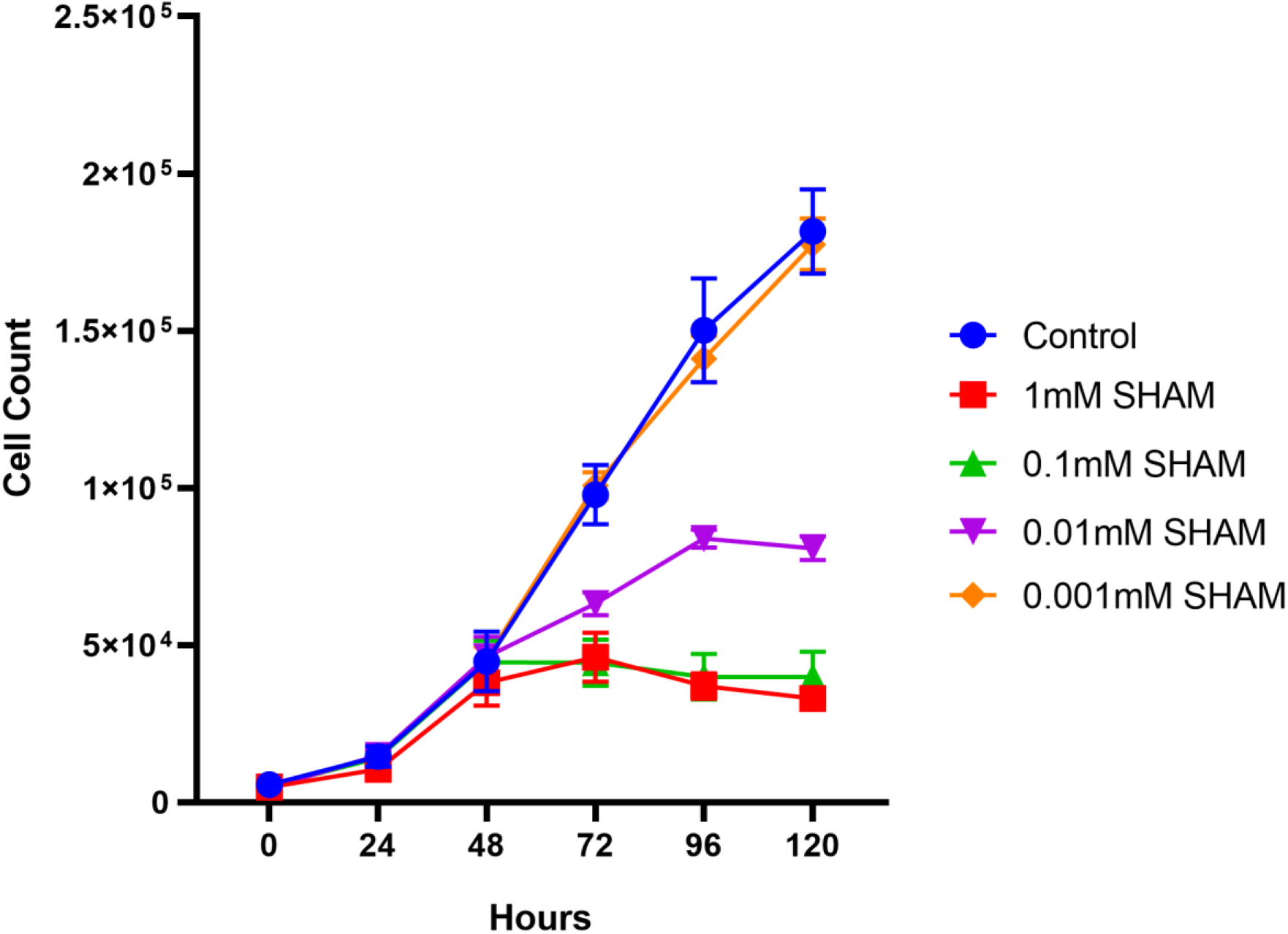
Growth of *N. gruberi* is reduced in increasing concentrations of SHAM. Using the JuLI^™^Stage Live cell monitoring system, we were able to monitor the growth rates of N. gruberi cells in varying concentrations of SHAM. Cells were seeded in wells of a 96-well plate and left to grow for 48 hours prior to the addition of SHAM at 1mM, 0.1mM, 0.01mM and 0.001mM concentration. By counting cells, we noticed a significant decrease in cell numbers when challenged with 1 mM and 0.1mM SHAM. In the presence of 0.01mM SHAM we saw a decrease in proliferation rate, peaking at 96 hours. We observed no effect using SHAM at 0.001mM. Error bars are standard error of the mean.

## DISCUSSION

The genome of *Naegleria gruberi* (Fritz-Laylin *et al*., 2010) encodes several homologues of alternative oxidases, but until now, there was no thorough report to investigate the origin and distribution of these proteins within microbial eukaryotes. Our sophisticated search and analyses suggest that the AOX gene was present in the ancestors of multiple eukaryotic groups, suggesting potential acquisition at earlier stages in the eukaryotic evolution. Specifically, for the *Naegleria* species, our phylogenetic analysis indicates that AOX gene duplication took place prior to speciation events, which further suggests the earlier requirement of two homologues as an adaptation to the unique lifestyles of these organisms. In *N. gruberi*, these homologues have been previously shown to be differentially expressed during the organism’s two major stages, trophozoite and flagellate (https://phycocosm.jgi.doe.gov/pages/blast-query.jsf?db=Naegr1) (Fritz-Laylin *et al*., 2010). The reason behind this is currently unknown and no conclusions can be extracted from the primary amino acid sequences of the two homologues. Our sequence alignment analysis reveals that the key residues required for AOX to be functional are present across all homologues and all three *Naegleria* species. One of the key domains presented in the sequences is the diiron domain, characterized by four glutamic acid residues and two histidine residues involved in forming hydrogen bonds with Fe^2+^ and Fe^1+^ ions, of which the latter are considered to be universally conserved (Moore *et al*., 2013). In addition, our sequence alignment reveals the presence of the three universally conserved tyrosine residues (Moore *et al*., 2013) in the electron transport chain (Moore *et al*., 2013; Shiba *et al*., 2013). Interestingly, our sequence alignment revealed a deviation in conservation of the residue trp-151 (*Tb*AOX), including between the pathogenic *N. fowleri*, non-pathogenic *N. lovaniensis* and, *N. gruberi* species. It has previously been reported that a mutation from trp-206 (trp-151 on tbAOX) to ala-206 resulted in a decrease in respiration of 95% (Crichton *et al*., 2010). However, wet lab experiments would be required to ascertain the effect of the difference in respiration found in *Naegleria* species.

Following up from the *in silico* analysis, we aimed to biochemically characterize the *N. gruberi* AOX homolog. It had been previously experimentally demonstrated that *N. gruberi*’s preferred food substrate are lipids over glucose, which was subsequently associated with the unique abundance of metabolites in the brain (Bexkens *et al*., 2018). The authors demonstrated that *N. gruberi* has a biochemically active AOX, but they did not show any localization. We have designed a peptide and generated an anti-*Naegleria* polyclonal antibody that could cross-react with all *Naegleria* species and be used for localization studies. Our immunoblotting results show a strong band appearing at 28 kDa, which was the predicted mass of *N. gruberi’*s AOX. This signal only appeared in the whole cell lysate and mitochondrial fraction, with no signal in the cytosolic fraction. This data was strengthened by our immunofluorescent microscopy experiments, which revealed a high degree of colocalization between the AOX staining and a commercial mitochondrial stain (Tsaousis *et al*., 2014). Lastly, an experiment using immunogold labelling showed staining signal from inside the mitochondria, which we find encouraging as AOX is located on the inner mitochondrial membrane (Berthold, Andersson and Nordlund, 2000).

Next, real-time respirometry data showed strong evidence of AOX being an active component in *N. gruberi*, since the addition of KCN did not lower the oxygen flux. Interestingly, previous work explored the potential cyanide resistance of *N. gruberi* as part of a larger metabolomic study (Bexkens *et al*., 2018). The authors reported an 80% decrease in respiration, which contrasts with our results. The authors also reported that addition of SHAM further decreased respiration by 14%. The differences between the results reported here and the aforementioned study is likely to be attributed to the differing growth conditions or prolonged differential adaptations of the laboratory strains (Schuster, 2002). Nonetheless, our cellular and biochemical data confirm their hypothesis regarding the presence of AOX in *N. gruberi* mitochondria. This however leads to significant questions in the pursuit of elucidating the complex metabolic mechanisms in *N. gruberi*, in particular, which environmental conditions favor expression and utilization of AOX, there is no report demonstrating the endurance of *Naegleria* species under, for example, hypoxic conditions. *Naegleria* species favor warm and moist environments and have been isolated from lakes, rivers, geothermal springs along with man-made bodies of water such as swimming pools, thermal effluent, sewage sludge and water-cooling circuits from power station [for review see (Chalmers, 2014)]. As a result, *Naegleria* species seem to acclimatize in various temperatures, turbidities and metal concentrations, which subsequently affect the mitochondrial functions. In other organisms, mainly plants, AOX has been implicated in metabolic and signaling hemostasis and was demonstrated to be particularly important during a variety of stresses, including alterations in temperature, nutrient deficiency, oxygen levels and metal toxicity (Vanlerberghe, 2013a; Saha, Borovskii and Panda, 2016). Such observations may be true for *Naegleria* as well, and further investigations using these different parameters are required to understand the true role of the various AOX homologs in *Naegleria*’s survival. For example, a previous report has shown differences between metabolic activities of iron-saturated and iron restricted trophozoites of *N. gruberi*, which could subsequently be attributed to potential iron homeostasis centrally regulated by the mitochondria (Mach *et al*., 2018). Again, it will be interesting to further investigate the role and function of the AOX in such mechanisms.

While reviewing the adaptations of *Naegleria* in the various environments, we cannot avoid discussing the potential role of AOX in the pathogenesis of *N. fowleri*. A recently published ‘omics approach investigating the potential genes that could be driving the pathogenicity of *N. fowleri* demonstrated up-regulation of mitochondrial energy conversion genes including those involved in ubiquinone biosynthesis, isocitrate dehydrogenase (TCA cycle), complex I and complex III of oxidative phosphorylation (Herman *et al*., 2020). While the authors have demonstrated up-regulation of enzymes indirectly involved in oxidative stress pathway (e.g. agmatine deiminase), they were not able to demonstrate any significant change in the expression levels of any of the *N. fowleri* AOX homologs (Herman *et al*., 2020). Based on these observations, it would be interesting to investigate the expression levels of AOX from the amoebas collected directly from the brain, either through transcriptomics and /or proteomics, as well as investigate whether this protein is essential for the survival of *N. fowleri* in such a complicated and variable environment.

It has been observed that there are significant differences in concentration levels of metabolites between various brain regions (Cichocka and Bereś, 2018). *N. fowleri* is typically found in olfactory bulb of the brain (Moseman, 2020), which it has a unique metabolic network signature and is highly abundant in various salts (e.g. sodium, potassium, calcium), metals (e.g. iron, copper and magnesium) (Gardner *et al*., 2017), metabolites (histidine-containing dipeptides such as anserine, carnosine, b-alanine), cholesterol, poly-unsaturated fatty acids and prostaglandins (Choi *et al*., 2018), as well as featuring variable concentrations of oxygen (Özugur, Kunz and Straka, 2020). Differences in the concentrations of these factors have been previously shown to provide stimuli for alterations in the expression of AOX in plants (Vanlerberghe, 2013) and trypanosomes (Vassella *et al*., 2004). As previously discussed, AOX was also shown to be implicated in metabolic and signaling hemostasis and was demonstrated to be particularly important during a variety of stresses, including alterations in temperature, nutrient deficiency, oxygen levels and metal toxicity (Saha, Borovskii and Panda, 2016). Additionally, it has been demonstrated that salt stress negatively impacts mitochondria function, resulting in decreased electron transport activities, with increased mitochondrial ROS and lipid peroxidation, followed by subsequent induction of mitochondrial ROS-scavenging systems, including increase activity of AOX (Ferreira *et al*., 2008; Mhadhbi *et al*., 2013; Saha, Borovskii and Panda, 2016). We speculate that similar implications could be associated with the function of AOX in *N. fowleri* populating the brain. Demonstrating that anti-AOX drugs are effective against *Naegleria* growth, is of great importance to investigate whether these proteins are essential for the survival of *N. fowleri* in the brain, and determine if it is possible to efficiently utilize these compounds (Murphy and Lang-Unnasch, 1999; Ebiloma *et al*., 2019; Barsottini *et al*., 2020) against the “brain-eating amoeba”.

Herein, we provide the first thorough investigation of the localization and functional characterization of the alternative oxidase proteins in *Naegleria* species. These single-protein-focused studies are essential in contributing to our understanding of the biochemical and cellular adaptations of this exceptional microbial eukaryote as well as provide another piece in *Naegleria*’s evolutionary puzzle. As a result, our investigation on the function of the AOX provides an additional step towards developing this organism as a model to understand various pan-eukaryotic adaptations and more importantly how metabolism could affect its opportunistic nature.

## ACKNOWLEDGEMENTS

This research was supported by BBSRC research grant (BB/M009971/1) to ADT. EK was supported by a Betty and Gordon Moore Foundation grant to ADT.

## AUTHORS CONTRIBUTIONS

D.C., A.O., S.G., C.W.G., and A.D.T. designed the experiments. D.C., A.O., N.T., E.K., I.R.B., and E.E. conducted the experiments. D.C., N.T. and G.T. performed the data analysis. D.C. and A.D.T. wrote the manuscript and all co-authors reviewed and approved it.

## Supplementary Figures

**Supplementary Figure S1. Sequence alignments of AOX reveals conserved domains**

Maximum likelihood tree of AOX homologs in eukaryotes and bacteria (323 sequences, 148 amino acid positions). The tree was inferred with IQTREE using the LG+I+G4 model selected under the BIC criterion. Grey dots correspond to supports higher than 80%. The scale bar corresponds to the average number of substitutions per site.

**Supplementary Figure S2. Sequence alignments of AOX reveals conserved domains**

To assess level of conservation between AOXs we aligned ngAOX, nfAOX and nlAOX amino acid sequences against other well characterized AOXs; *Trypanosoma brucei, Candida albicans, Arabidopsis thaliana, Cryptosporidium parvum* and, *Neurospora crassa*. We observed a considerable amount of conservation between all AOXs towards the middle and C-terminal end, where the presence of the alpha helical bundles and di-iron binding domains reside. * denotes key amino acids presented in Table below. † denotes a deviation in conserved amino acids of AOX

**Supplementary Figure S3. Confocal microscopy demonstrating mitochondrial localization of AOX.**

Additional *N. gruberi* cells demonstrating localization of AOX in their mitochondria. *N. gruberi* cells were treated with mitotracker red prior to fixation and then probed with AOX antibodies (green). Nuclear marker and mtDNA staining is shown in blue (DAPI staining) Our confocal imaging reveals a high degree of co-localization between the mitotracker red signal and the green signal derived from immunoprobing AOX.

## Notes

### Competing Interest Statement

The authors have declared no competing interest.

