## Supplementary figures and images for "Localization and Functional Characterization of the Alternative Oxidase in *Naegleria*"

### Supplementary Figure 1

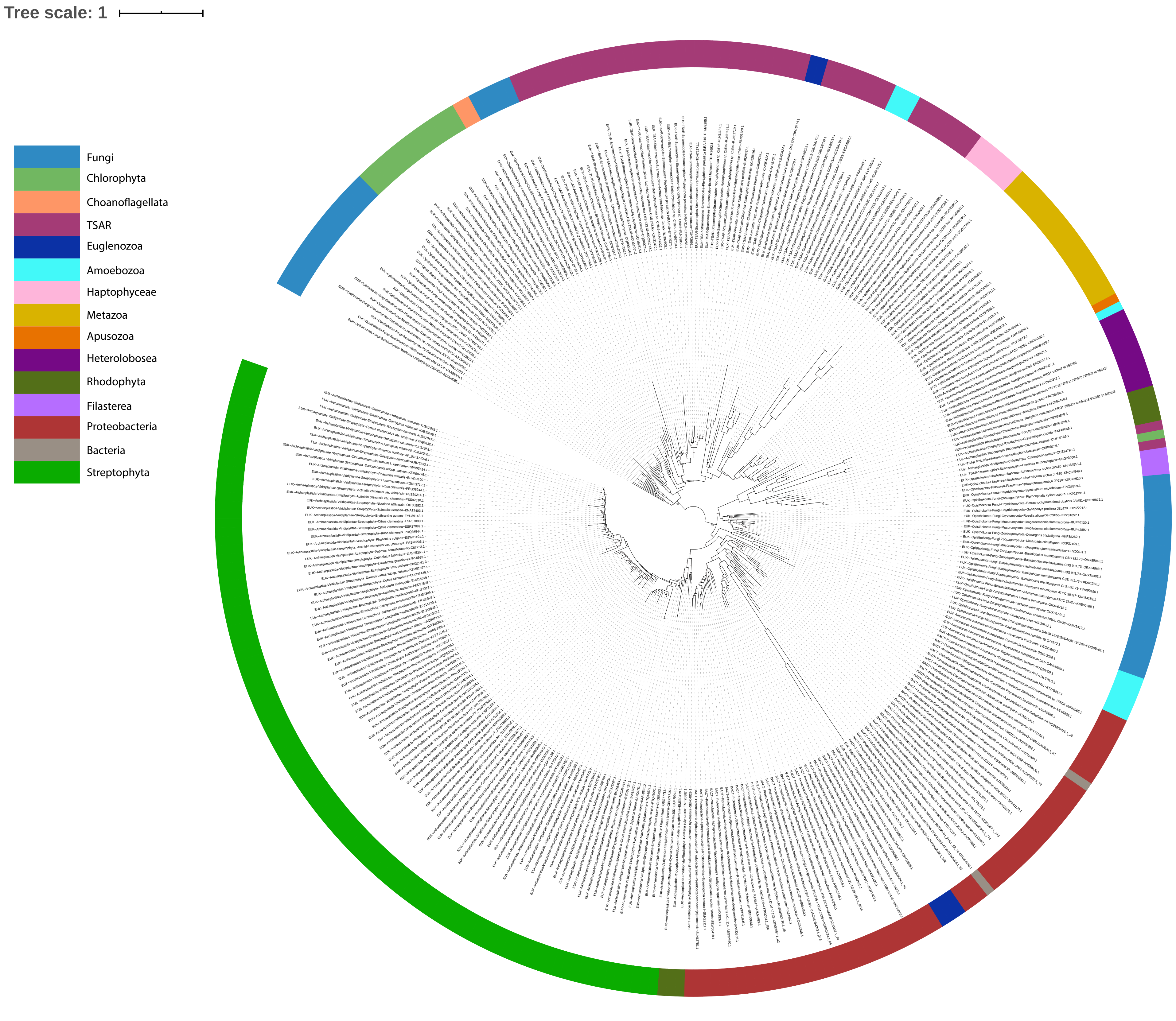

### Supplementary Figure 3

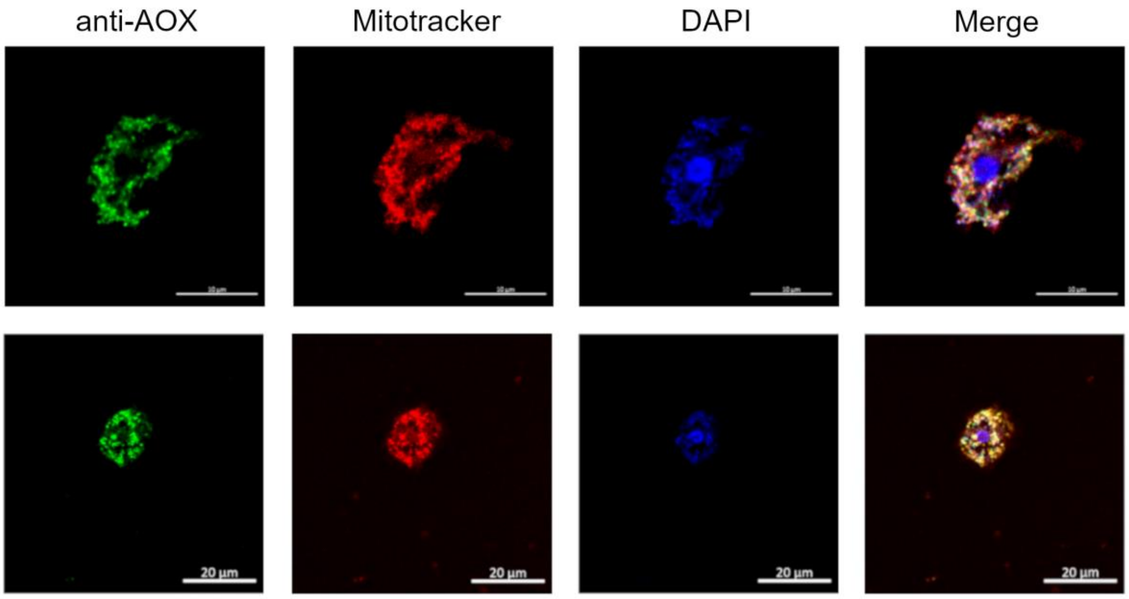
