## Supplementary Figure 2 for "Localization and Functional Characterization of the Alternative Oxidase in *Naegleria*"

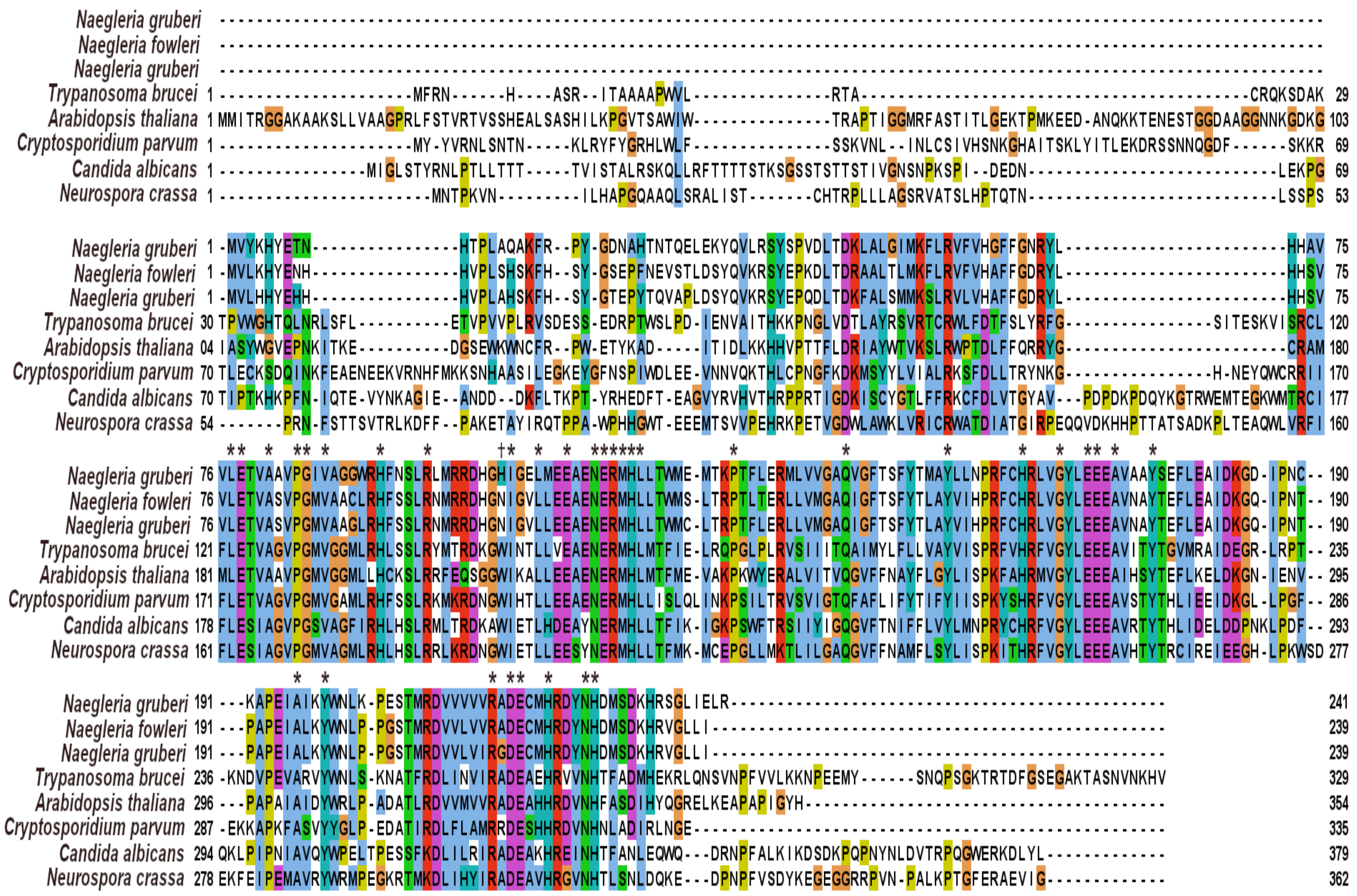
Naegleria gruberi
Naegleria fowleri
Naegleria gruberi
Trypanosoma brucei 1 --------------------MFRN-----H---ASR--ITAAAAPWVL--------------RTA-----------------------------------CRQKSDAK 29
Arabidopsis thaliana 1 MMITRGGAKAAKSLLVAAGPRLFSTVRTVSSHEALSASHILKPGVTSAWIW--------------TRAPTIGGMRFASTITLGEKTPMKEED-ANQKKTENESTGGDAAGGNNKGDKG 103
Cryptosporidium parvum 1 --------------------MY-YVRNLSNTN----KLRYFYGRHLWLF--------------SSKVNL--INLCSIVHSNKGHAITSKLYITLEKDRSSNNQGDF-----SKKR 69
Candida albicans 1 ---------------MIGLSTYRNLPTLLTTT----TVISTALRSKQLLRFTTTTSTKSGSSTSTTSTIVGNSNPKSPI--DEDN--------------------------LEKPG 69
Neurospora crassa 1 ---------------------MNTPKVN---------ILHAPGQAAQLSRALIST------CHTRPLLLAGSRVATSLHPTQTN---------------------------LSSPS 53
Naegleria gruberi 1 -MVYKHYETN---------------HTPLAQAKFR--PY-GDNAHTNTQELEKYQVLRSYSPVDLTDKLALGIMKFLRVFVHGFFGNRYL---------------------------HHAV 75
Naegleria fowleri 1 -MVLKHYENH---------------HVPLSHSKFH--SY-GSEPFNEVSTLDSYQVKRSYEPKDLTDRAALTLMKFLRVFVHAFFGDRYL---------------------------HHSV 75
Naegleria gruberi 1 -MVLHHYEHH---------------HVPLAHSKFH--SY-GTEPYTQVAPLDSYQVKRSYEPQDLTDKFALSMMKSLRVLVHAFFGDRYL---------------------------HHSV 75
Trypanosoma brucei 30 TPWWGHTQLNRLSFL----------ETVPVVPLRVSDESS-EDRPTWSLPD-IENVAITHKKPNGLVDTLAYRSVRTCRWLFDTFSLYRFG--------------SITESKVISRCL 120
Arabidopsis thaliana 04 IASYWGVEPNKITKE----------DGSEWKWNCFR--PW-ETYKAD-----ITIDLKKHHVPTTFLDRIAYWTVKSLRWPTDLFFQRRYG------------------------CRAM 180
Cryptosporidium parvum 70 TLECKSDQINKFEAENEEKVRNHFMKKSNHAASILEGKEYGFNSPIWDLEE-VNNVQKTHLCPNGFKDKMSYYLVIALRKSFDLLTRYNKG--------------H-NEYQWCRRII 170
Candida albicans 70 TIPTKHKPFN-IQTE-VYNKAGIE--ANDD--DKFLTKPT-YRHEDFT-EAGVYRVHVTHRPPRTIGDKISCYGTLFFRKCFDLVTGYAV---PDPDKPDQYKGTRWEMTEGKWMTRCI 177
Neurospora crassa 54 -------PRN-FSTTSVTRLKDFF--PAKETAYIRQTPPA-WPHHGWT-EEEMTSVVPEHRKPETVGDWLAWKLVRICRWATDIATGIRPEQQVDKHHPTTATSADKPLTEAQWLVRFI 160
** * ** * * * †* * * ****** * * * * * ** * *
Naegleria gruberi 76 VLETVAAVPGIVAGGWRHFNSLRLMRRDHGHIGELMEEAENERMHLLTWME-MTKPTFLERMLVVGAQVGFTSFYTMAYLLNPRFCHRLVGYLEEEAVAAYSEFLEAIDKGD-IPNC-- 190
Naegleria fowleri 76 VLETVASVPGMVAACLRHFSSLRNMRRDHGNIGVLLEEAENERMHLLTWMS-LTRPTLTERLLVMGAQIGFTSFYTLAYVIHPRFCHRLVGYLEEEAVNAYTEFLEAIDKGQ-IPNT-- 190
Naegleria gruberi 76 VLETVASVPGMVAAGLRHFSSLRNMRRDHGNIGVLLEEAENERMHLLTWMC-LTRPTFLERLLVMGAQIGFTSFYTLAYVIHPRFCHRLVGYLEEEAVNAYTEFLEAIDKGQ-IPNT-- 190
Trypanosoma brucei 121 FLETVAGVPGMVGGMLRHLSSLRYMTRDKGWINTLLVEAENERMHLMTFIE-LRQPGLPLRVSIIITQAIMYLFLLVAYVISPRFVHRFVGYLEEEAVITYTGVMRAIDEGR-LRPT-- 235
Arabidopsis thaliana 181 MLETVAAVPGMVGGMLLHCKSLRRFEQSGGWIKALLEEAENERMHLMTFME-VAKPKWYERALVITVQGVFFNAYFLGYLISPKFAHRMVGYLEEEAIHSYTEFLKELDKGN-IENV-- 295
Cryptosporidium parvum 171 FLETVAGVPGMVGAMLRHFSSLRKMKRDNGWIHTLLEEAENERMHLLISLQLINKPSILTRVSVIGTQFAFLIFYTIFYIISPKYSHRFVGYLEEEAVSTYTHLIEEIDKGL-LPGF-- 286
Candida albicans 178 FLESIAGVPGSVAGFIRHLHSLRMLTRDKAWIETLHDEAYNERMHLLTFIK-IGKPSWFTRSIIYIGQGVFTNIFFLVYLMNPRYCHRFVGYLEEEAVRTYTHLIDELDDPNKLPDF-- 293
Neurospora crassa 161 FLESIAGVPGMVAGMLRHLHSLRRLKRDNGWIETLLEESYNERMHLLTFMK-MCEPGLLMKTLILGAQGVFFNAMFLSYLISPKITHRFVGYLEEEAVHTYTRCIREIEEGH-LPKWSD 277
* * * ** * **
Naegleria gruberi 191 ---KAPEIAIKYWNLK-PESTMRDVVVVVRADECMHRDYNHDMSDKHRSGLIELR------------------------------------------ 241
Naegleria fowleri 191 ---PAPEIALKYWNLP-PGSTMRDVVLVVRADECMHRDYNHDMSDKHRVGLLI-------------------------------------------- 239
Naegleria gruberi 191 ---PAPEIALKYWNLP-PGSTMRDVVLVIRGDECMHRDYNHDMSDKHRVGLLI-------------------------------------------- 239
Trypanosoma brucei 236 -KNDVPEVARVYWNLS-KNATFRDLINVIRADEAEHRVVNHTFADMHEKRLQNSVNPFVVLKKNPEEMY------SNQPSGKTRTDFGSEGAKTASNVNKHV 329
Arabidopsis thaliana 296 ---PAPAIAIDYWRLP-ADATLRDVVMVVRADEAHHRDVNHFASDIHYQGRELKEAPAPIGYH---------------------------------- 354
Cryptosporidium parvum 287 -EKKAPKFASVYYGLP-EDATIRDLFLAMRRDESHHRDVNHNLADIRLNGE---------------------------------------------- 335
Candida albicans 294 QKLPIPNIAVQYWPELTPESSFKDLILRIRADEAKHREINHTFANLEQWQ---DRNPFALKIKDSDKPQPNYNLDVTRPQGWERKDLYL----------- 379
Neurospora crassa 278 EKFEIPEMAVRYWRMPEGKRTMKDLIHYIRADEAVHRGVNHTLSNLDQKE---DPNPFVSDYKEGEGGRRPVN-PALKPTGFERAEVIG------------ 362

Table 1. Universally conserved amino acids of AOX and their roles. Comparison of amino acid residues between *Trypanosomas brucei brucei* AOX (trAOX) and *Naegleria gruberi* AOX (ngAOX), *Naegleria fowleri* AOX (nfAOX) and, *Naegleria lovaniensis* AOX (nlAOX)

| **tbAOX numbering** | **ngAOX numbering** | **nfAOX numbering** | **nlAOX numbering** | **Role** |
|---|---|---|---|---|
| Leu-122 | Leu-77 | Leu-77 | Leu-77 | Substrate-binding channels 1 and 2 |
| Glu-123[†] | Glu-78 | Glu-78 | Glu-78 | Fe-Fe Ligand; substrate binding channels 1 and 2 |
| Ala-126 | Ala-81 | Ala-81 | Ala-81 | Substrate-binding channels 1 and 2 |
| Pro-129 | Pro-84 | Pro-84 | Pro-84 | Substrate-binding channels 2 |
| Gly-130 | Gly-85 | Gly-85 | Gly-85 | Forms kink in helix α2 |
| Val-132 | Val-87 | Val-87 | Val-87 | Hydrophobic interaction with helix α6 |
| His-138 | His-93 | His-93 | His-93 | Membrane-binding region/dimer interface |
| Arg-143 | Arg-98 | Arg-98 | Arg-98 | Membrane-binding region/dimer interface |
| Trp-151 | *His-106* | *Asn-106* | *Asn-106* | *Hydrophobic interaction with helix α6* |
| Iso-152 | Iso-107 | Iso-107 | Iso-107 | Dimer interface |
| Leu-155 | Leu-110 | Leu-110 | Leu-110 | Dimer interface |
| Glu-158 | Glu-113 | Glu-113 | Glu-113 | Substrate-binding channels 1 and 2 |
| Asn-161[†] | Asn-116 | Asn-116 | Asn-116 | Secondary ligation sphere; hydrogen bond network |
| Glu-162[†] | Glu-117 | Glu-117 | Glu-117 | Fe-Fe Ligand |
| Arg-163 | Arg-118 | Arg-118 | Arg-118 | Membrane-binding region |
| Met-164 | Met-119 | Met-119 | Met-119 | Dimer interface; interaction with N-terminal arm |
| His-165 | His-120 | His-120 | His-120 | Fe-Fe ligand |
| Leu-166 | Leu-121 | Leu-121 | Leu-121 | Dimer interface |
| Pro-175 | Pro-130 | Pro-130 | Pro-130 | Dimer interface |
| Gln-187 | Gln-142 | Gln-142 | Gln-142 | Dimer interface |
| Tyr-198[†] | Tyr-153 | Tyr-153 | Tyr-153 | Dimer interface; hydrogen bonds to His-206 (tbAOX) |
| His-206 | His-161 | His-161 | His-161 | Membrane-binding region |
| Gly-210 | Gly-165 | Gly-165 | Gly-165 | Forms kink in helix α5 |
| Glu-213[†] | Glu-168 | Glu-168 | Glu-168 | Fe-Fe ligand |
| Glu-214 | Glu-169 | Glu-169 | Glu-169 | Interacts with N-terminal arm |
| Ala-216 | Ala-171 | Ala-171 | Ala-171 | Substrate-binding channel 2 |
| Tyr-220 [†] | Tyr-175 | Tyr-175[†] | Tyr-175 | Catalytic cycle |
| Ala-243 | Ala-196 | Ala-196 | Ala-196 | Hydrophobic interaction with helix α3 |
| Tyr-246[†] | Tyr-199 | Tyr-199 | Tyr-199 | Secondary ligation sphere; hydrogen bond network |
| Arg-263 | Arg-216 | Arg-216 | Arg-216 | Interaction with helix α5 and N-terminal arm |
| Asp-265[†] | Asp-218 | Asp-218 | Asp-218 | Secondary ligation sphere; hydrogen bond network |
| Glu-266 | Glu-219 | Glu-219 | Glu-219 | Fe-Fe ligand |
| His-269 | His-222 | His-222 | His-222 | Fe-Fe ligand |
| Asn-273 | Asn-226 | Asn-226 | Asn-226 | Interaction with helix α5 |
| His-274 | His-227 | His-227 | His-227 | Interaction with helix α5 |

[†]Denotes universally conserved residues of AOX
